# Intermittent compression increases flow through a gel: a mechanical model perhaps relevant to flow of cerebrospinal fluid

**DOI:** 10.1101/2021.12.30.474515

**Authors:** Michael G. Hale, Jonathan A. Coles

## Abstract

Exchange of molecules between cerebrospinal fluid (CSF) and brain cells contributes to brain function and protection from dementia. Despite widespread acceptance of the ‘glymphatic’ theory, the route by which CSF is brought close enough to the neural tissue for solutes to be exchanged by extracellular diffusion is not entirely clear. Exogenous molecules injected into CSF are observed to reach the basement lamina that surrounds the dense capillary network. Transport of solutes by diffusion along the basement lamina, a gel of macromolcules about 100 nm thick, would be too slow; bulk flow in a static geometry would require unphysiologically high pressures. However, it is known that the pulsation of blood aids transport of CSF, and we hypothesized that this is because the pulsation intermittently squeezes the pericapillary lamina. In a primitive mimicry, we have tested whether intermittent squeezing increases flow through an agar gel. In all but one of 216 tests, pulsation caused a reversible increase, sometimes by a factor of 100 or more. The enhancement was greatest for frequencies 5-11 Hz and, over the tested range of pressure heads (20 - 50 cmH_2_O), was greatest for the lowest pressure. The results suggest a reason why exercise slows the aging of the brain.

## Introduction

Blood supplies the main molecules required for brain metabolism, glucose and oxygen. Arterioles penetrating into the brain from the surface carry blood to a dense capillary network drained by venules that return it to veins on the brain surface (Fig 1A,C). Glucose and oxygen can escape across the capillary wall and then diffuse the short distances (on average 13 μm, [1, 2]) to the neurons and glial cells. A second, complementary, source of other necessary molecules, including vitamins and DNA precursors, is cerebrospinal fluid (CSF) (reviewed by [3, 4]). CSF can also carry away molecules, such as amyloid beta, whose accumulation in the brain is associated with Alzheimer’s disease [5]. CSF is secreted into the brain ventricles by the choroid plexuses which synthesize, or transport from blood, a palette of molecules apparently necessary to brain maintenance. By injecting marker molecules visible in microscopy it has been shown that CSF flows through ducts connecting the ventricles to the surface of the brain at the cisterna magna and from there is distributed to the surface of the cortex [6–12]. It is widely agreed that CSF next flows down a perivascular space of the arterioles that leave the cortical surface and penetrate into the neural tissue, that CSF somehow goes from there to the perivascular spaces of the venules and that it then returns to the meninges and is removed from the cranium (Fig. 1C [4]). There are two main theories about how the CSF (or a derived fluid) passes from the peri-arteriolar space to the perivenular space. Currently the most popular is the ‘glymphatic’ theory, originally based on a landmark imaging paper by Iliff et al. [9]. The glymphatic theory [9, 13] states that CSF crosses the layer of astrocyte endfeet that delimit the peri-arteriolar space and moves by bulk (convective) flow through the extracellular matrix of the neural tissue to reach the perivascular space of venules. An alternative, earlier theory, which we will call the ‘paracapillary’ theory, is that CSF flows along paravascular conduits that accompany the capillaries that link the arterioles to the venules [7, 14–17] [18]

**Fig 1.**
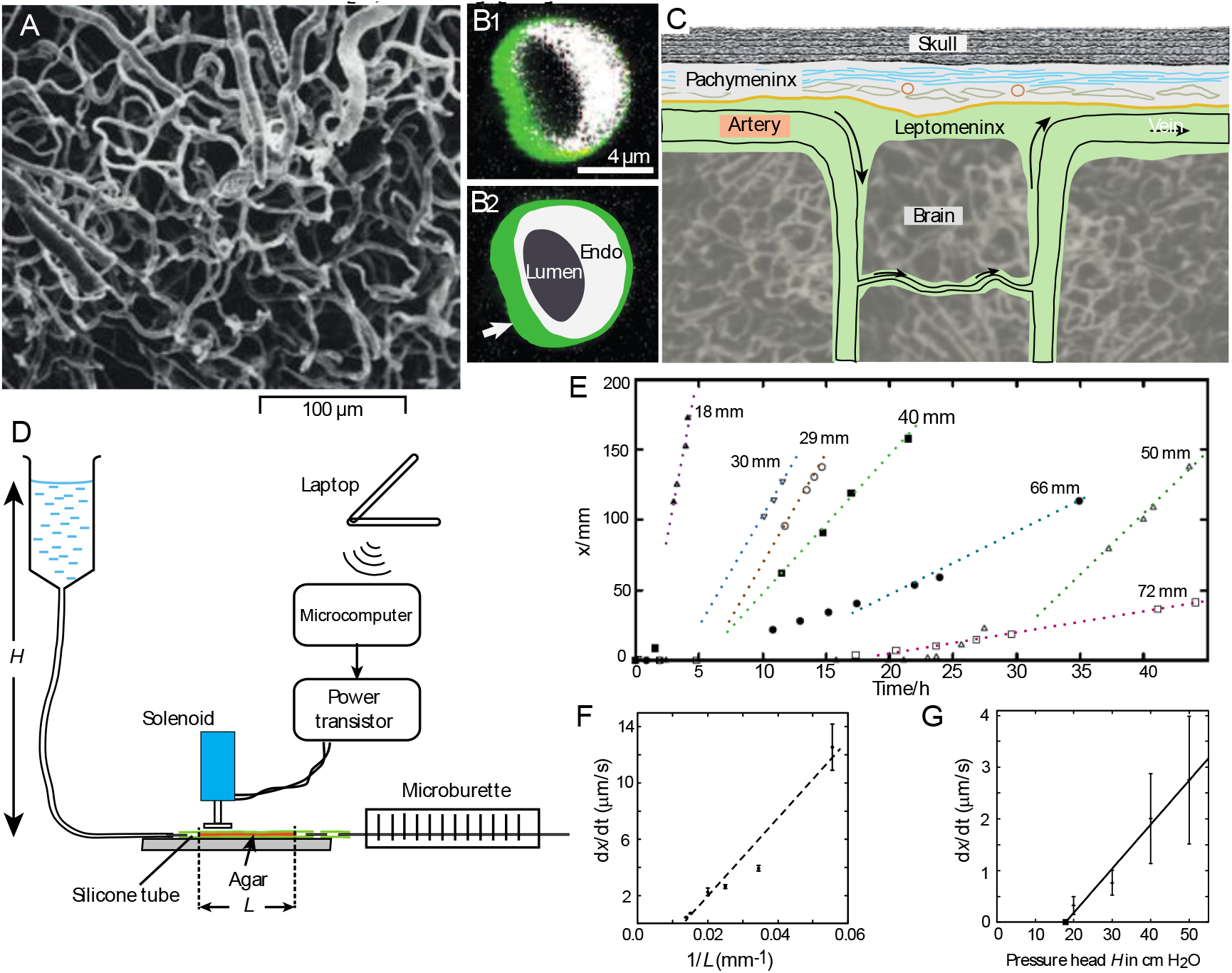
Flow without pulsation. **A.** A cast of the blood vessels in the cortex of a rat brain (modified from [2] with permission from the Journal of Neurosurgery). **B_1_** A CSF marker dye in a space surrounding a capillary in a section of fixed mouse brain. Endothelium is labelled by expression of GFP (shown in white) driven by the Tie-2 promoter. Ovalbumin conjugated with Alexa Fluor 647 (M.Wt. 45 kDa, shown as green) was injected in the cisterna magna less than 30 min before perfusion fixation (from Fig. 3C in [9] Reprinted with permission from AAAS). **B_2_** Interpretation of **B_1_** indicating the lumen of the capillary, its endothelial cell wall and (arrow) the basement lamina. **C.** Scheme of the pathway of CSF in the cortex proposed by [7, 14, 16, 17, 19] showing flow of CFS along a pericapillary space. CSF flows in the subarachnoid space of the leptomeninx, the inner of the two meningeal compartments, which is separated from the outer compartment, the pachymeninx, which contains a layer of collagen (blue), the dura mater. **D.** Minimal portrayal of the experimental set-up (see Supp Mat 1 for details). **E.** Displacement, *x*, of the bubble in the microburette caused by flow through gel columns of different lengths *L*, with a static pressure gradient of 40 cm H_2_O. The dotted lines (drawn by eye) show that in most cases the flow took time to develop. **F.** The mean flow rates for the linear parts of **E** plotted against 1/*L*. **G**. Flow rates through columns approximately 40 mm long at different pressure heads *H*. Bars are SEMs. The regression line through the data points does not extrapolate to zero.

Strong objections have been made to both theories. One objection to the glymphatic theory is that bulk flow through the extracellular matrix would require unphysiologically high pressure gradients [12, 20–22]. Another apparent weak point is that it has repeatedly been observed that molecules carried by CSF are found, concentrated relative to adjacent tissue, in the basement lamina that surrounds brain capillaries (Fig 1B; [7, 15, 23] [9, 10]. The great objection to the paracapillary theory is that there is no obvious paracapillary conduit, the extracellular space between the capillary endothelium and the neural tissue being occupied by macromolecular matrices, notably the basement lamina [17, 24]. It has also been suggested that the observed labelling of the basement lamina is an artifact: since current intravital imaging techniques lack the resolution needed to clearly image the capillary basement lamina (which is about 100 nm thick) these observations were made on tissue chemically fixed after death. It has been proposed that CSF reaches this location only after cardiac arrest, perhaps because of the sudden drop in blood pressure [25–27]. However, control images of living capillaries are lacking, and the experimental evidence of Rennels *et al*. [7] is the opposite: they found that aortic occlusion reduced the transport of CSF marker.

The key objection to both theories, that an unphysiologically high pressure gradient would be required to drive bulk flow, supposes that the microstructure of the fluid path is static. Experiments have shown that delivery of CSF to pericapillary space in the brain (and also maintainance of brain function) is greatly reduced if the pulsation of the circulating blood is reduced [7, 28]. Analogous results have been found for bulk flow through the extracellular matrix of rabbit skin [29]. Blood pulsation also appears to assist CSF flow along the paravascular spaces of cerebral arteries and arterioles [9, 10, 27, 30] and, at least in the dorsal cortex, the venules pulsate with an amplitude even greater than that of the descending arterioles [10]. Hence the blood pressure in the capillaries must also pulsate. Although, by Laplace’s Law, small tubes such as capillaries are resistant to pressure changes, we cannot exclude changes in diameter sufficient to cause significant changes in pressure in the immediate vicinity of the capillary.

There is an extensive literature on theories of flow through porous media (see e.g., [31]) particularly concerning the permeability of soil and gravel [32–34] and carbon microstructures [35, 36]. The relatively rare experimental studies confirm that resistance to flow through carbon nanotubes can be very small [37] and that flow through soil or porous rock is not simply proportional to the pressure gradient [38–42]. However, we find no report of experiments on the effect on flow through a matrix of macromolecules in a defined geometry of applying pulsating lateral pressure changes. The closest seems to be the experiment of Parsons and McMasters [43] in which they applied intermittent pressure to rabbit skin and found that it speeded the spread of extracellular dye. It is not trivial to make a laboratory model that mimics the pericapillary space and apply a pulsating pressure, uniform along its length. So we have done a ‘proof-of-principle’ experiment just to see if pulsation affects flow through a gel contained in an elastic tube. We found that pulsation markedly and reversibly increased the flow, sometimes by a factor of a hundred or more. We have examined the dependence of the flow on the degree of compression of the tube, the frequency of pulsation, and the static pressure head. We conclude that in the presence of a pulsating blood flow, bulk paracapillary flow of CSF is a possibility that merits further investigation.

## Methods

### Preparation of gel columns and the apparatus for pulsation

Silicone tubing, i.d. 1.0 mm, o.d. 3.0 mm, was cut into 40 - 80 mm lengths, filled with agar agar (Falooda powder, Top op Foods Ltd, UK), 1% w/v, prepared, and stored in, 0.15M NaCl. Five mm of gel was flushed from each end of a length of tube and it was connected upstream to a pressure head of 20 - 50 cm H_2_O and downstream to a microburette consisting of polyethylene tubing, i.d. 0.38 mm, held against a 200 mm scale (Fig 1D)A small index bubble was introduced into the microburette so that the movement of fluid could be measured. The gel-filled tube was held under a solenoid-driven piston, diameter 3.5 mm, that could pummel the tube at a frequency controlled by a microcomputer (Raspberry Pi zero-W) instructed by wifi from a laptop (SM2). The solenoid was mounted on a micromanipulator with a Vernier scale so that the approximate compression of the silicone tube could be controlled. Experiments were done at 18-22 °C. More details of the Methods are given in Supplementary Material 1.

### Control Experiment: Periodic compression slowed the flow of water through a fluid-filled elastic tube

Water with a head of about 30 cm flowed through a horizontal silicone tube, i.d. 1.0 mm, which passed beneath a small solenoid (Fig.SM2A). The flow was adjusted with a control valve to about 0.1 ml/sec. The water was switched to a coloured solution (CuSO_4_) and changes in the optical density at a point downstream of the solenoid were monitored using a red LED and a photodiode (Fig. SM2B). Squeezing the tube at 2 - 5 Hz with the solenoid slightly reduced the flow, as might be expected as the lumen of the tube was constricted for part of the time (Fig. SM2C). Squeezing produced no marked difference in the time course of the progressive increase in optical density caused by the arrival of the solute.

## Results

### Flow through the gel without pulsation

The volume flow, *Q*, through the column of 1% agar was measured by the displacement *x* of the bubble along the microburette scale. Results for six gel columns with lengths *L* = 18 - 72 mm at a pressure head *H* = 400 mm H_2_O are shown in Fig. 1E. The speed of displacement, d*x*/d*t*, was approximately constant for most of the observation period. There was often an initial period during which the bubble moved very slowly, so that extrapolation from the linear phase did not pass through the origin (dashed lines in Fig. 1E). When gel columns of different lengths were compared, the expected relationship d*x*/d*t* ∝ 1/*L* [32] was approximately observed (Fig. 1F). When the pressure head *H* was varied between 20 and 50 cm H_2_O, flow rate increased with *H* at pressures beyond a threshold of about 182 mm H_2_O (Fig. 1G). Such deviation from Darcy’s Law at low pressures is an accepted feature of flow through porous mineral beds [38, 42, 44, 45]. We calculated Darcy’s permeability coefficient (the mean specific hydraulic conductivity, see [46])

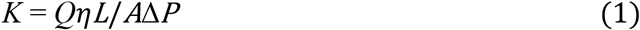

where *Q* is the volume flow, *η* is the viscosity of the fluid, taken as 1.06 mPa.sec [47], *L* is the length of the gel column, *A* is the cross-section of the silicone tube and *ΔP* is the pressure difference across the gel. *ΔP* = *ρg H* where *ρ* is the density of the fluid, and *H* is the height of the reservoir above the gel. The data shown in Fig. 1G give a range of *K* of 3.9-10.1 × 10^−16^ m^2^, with a mean of 7.6 × 10^−16^ m^2^. The SDs of individual measurements are in the order of the mean values.

### Pulsation increased the flow

In all but one of more than 216 tests, periodic compression (‘pulsation’) of the silicone tube, with a duty cycle of 50%, reversibly increased the flow of water through the column of agar gel. Results from a typical experiment are shown in Fig 2A: in this experiment, pulsation at 1 - 15 Hz increased the flow by factors of 3 - 11. For small compression depths, up to 0.5 mm, the enhancement increased monotonically with compression depth (Fig 2B); at greater compression depths no systematic trend was observed in the noise (not shown). If the pulsation was applied when the speed of the index bubble was very small (< 1 mm/h, see Fig 1C), the apparent enhancement could be very great: values over one hundred were excluded from averages.

**Fig 2.**
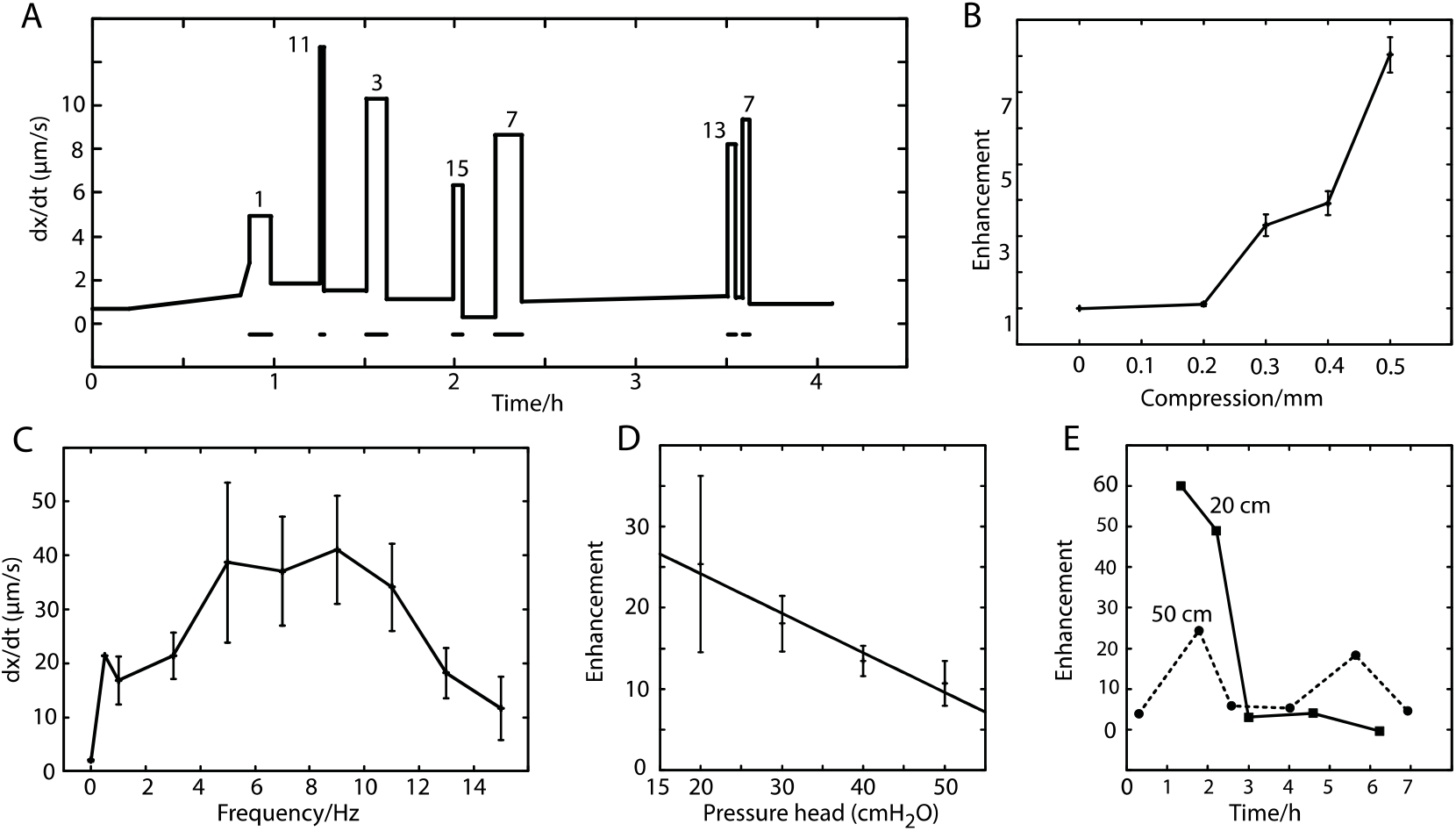
Pulsation increased flow through the gel. **A**. Results from one gel column. The average speed between readings of the index bubble in the microburette is plotted against time. Bars indicate pulsation at the frequencies given in Hertz. *L* = 16 mm, *H* = 30 cm. **B**. Enhancement of flow by pulsation at 2 Hz *vs* the compression by the solenoid. The data are for one gel column 30 mm long and a pressure head of 30 cm H_2_O. One outlying data point (much greater enhancement) was excluded **C**. Mean speed in the microburette as a function of pulsation frequency. Results on 10 gel columns with *L* = 38-41 mm, *H* = 40 cm. 6 - 15 measurements per frequency (except for 0.5 Hz). Bars are SEMs: there was great variation between measurements. With no pulsation the flow rate was 2.03 ± 0.25 μm/s. **D**. Mean flow enhancement vs pressure head at 5-11 Hz. Pooled results from a total of 55 measurement cycles with *L* = 16 - 53 mm, *f* = 5 - 11 Hz. **E**. Result from a single experiment in which *H* was alternated between 20 cm and 50 cm. Enhancement changed radically over tens of minutes. *L* = 30 mm, *f* = 7 Hz.

### Dependence on frequency

Fig 2C shows pooled results from ten gel columns of the speed of displacement of the index bubble. Bouts of pulsation at different frequencies were applied in pseudo-random order. Dividing *Q*(*f*) by the mean of the baselines, *Q*(0), just before and just after the pulsation, did not give a cleaner result, partly because in some cases the flow without pulsation was very small or undetectable. The maximum flow was for frequencies in the range 5 - 11 Hz.

### Dependence on pressure head

Pooled results from 55 tests showed that with pressure heads ranging from 20 cm H_2_O to 50 cm, the mean enhancement was greatest with *H* = 20 cm and fell as the pressure head was increased to 50 cm H_2_O (*P* = 0.022, Fig 2D).

### Variability

Despite the trend in the pooled results of Fig 2D, in individual experiments great variability was observed. For example, in the extreme case shown in Fig 2E the enhancement with *H* = 20 cm H_2_O was greater than that for 50 cm for some hours but switched to being less after 2.9h.

## Discussion

### Characteristics of flow through agar without pulsation

We have found no previous measurements of the permeability of 1% agar. Our measurements gave a mean Darcy coefficient of 7.6 × 10^−16^ m^2^ which is close to the value of 6.16 × 10^−16^ m^2^ found for 2% agarose by [48], but 579 times less than that of [49] (4.4 × 10^−13^ m^2^). The plot of bubble speed vs pressure head (Fig 1G) shows that there was little or no flow at the lowest pressures, in qualitative agreement with results on flow through soil and fractured rock [38, 42, 44, 45]. This behavior has been attributed to increased viscosity at the solid-liquid interface [50]. Previous experimenters have reported that flow could be initially very slow then increase after times in the order of minutes [41, 48]. In qualitative agreement with this, we also observed delays which, at the lowest pressure gradients, could last for several hours (Fig 1E).

### The effects of pulsation

We have found no previous report of the effect of applied pulsation on flow through a gel (apart from experiments on skin [43]). The increase in flow was nearly always reversed when the pulsation stopped, as in Fig 2A, which is evidence that the structure of the gel was not permanently changed. (In the cases where the flow did not reverse completely, a possibility is that gel was torn from the tube wall.) Presumably, beneath the solenoid piston, the main processes were a temporary distortion of the matrix as the vertical diameter of the tube was reduced and its width increased, and a temporary driving out of saline, upstream and downstream. Several modeling studies on flow through porous materials have concluded that water can become structured and more viscous at the water-solid interface [51, 52] and the experimental observation that K often decreases when the fluid pressure gradient is increased suggests that this structure can be disrupted when shear forces are increased [38, 41, 42]. In the present experiments the distortion of the gel will have increased shear forces at the water-matrix interface and this may have reduced the vicosity, allowing the greater bulk flow that was observed.

Since the area of fluid-solid interface, both within our gel model and in pericapillary tissue, is almost entirely within matrices of macromolecules rather than at macroscopic solid surfaces, it is to be expected that the effects of pulsation will be little affected by the macroscopic geometry. Therefore, it is perhaps more than a fortuitous coincidence that the averaged experimental data showed that maximum flow was produced by pulsation in the range 3 - 13 Hz which overlaps the range of heart rates in mice (6.6 - 13 Hz [53]) and is close to the range in humans (1 - 3 Hz).

Averaged data also showed a significant tendency for pulsation-induced enhancement of bulk flow to decrease as the pressure head was increased over the tested range of 20 - 50 cmH_2_O applied to a 40 mm column (Fig. 2D). This is to be expected as at low pressure heads the boundary layer will be thicker and its disruption by pulsation will have more effect on bulk flow.

### Relevance to a hypothetical flow along the pericapillary basement lamina

The pulsation of the penetrating arterioles [10] will produce pulsatile pressures on the capillary basement lamina at its proximal end. Venules downstream of brain capillaries are also observed to pulsate [10] indicating that the blood flow through the capillaries that feed into them is pulsatile. Unless the capillary endothelium is perfectly inelastic, this must result in some pulsatile compression of the basement lamina. If the hydraulic conductivity of the pulsed lamina were similar to that of 1% agar, could it support transport of physiological fluxes of CSF under physiological pressure gradients? The greater the CSF flux, the greater the forces needed to drive it so we conservatively consider the maximum possible CSF flux. Since it is not known how much of the CSF that reaches the leptomeninx then irrigates the neural tissue [4], we consider the extreme hypothesis that all CSF produced by the choroid plexuses is distributed to flow along paracapillary conduits. The rate of production of CSF in mouse is disputed: two reports give 0.33 - 0.37 μL/min [54, 55] whereas [56] give 0.1 μL/min. The volume of the mouse brain is 0.43 cm^3^ [57] so a high estimate of the production per unit volume is 0.86 μL min^−1^ cm^−3^, which is similar to that in the rat [58–61]. The total length of capillaries per unit volume of brain (in mouse, rat and cat) is about 1000 mm/mm^3^ [1, 62–64] and the mean length of individual capillaries is about 100 μm [2]. Hence, there are 10^4^ capillaries per mm^3^ or 10^7^ per cm^3^ and the paracapillary flow of CSF along each capillary is

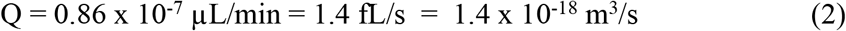

Taking the external diameter of a brain capillary as 4 μm [1, 62, 65] and the thickness of the basement lamina as 0.1 μm [66–68], the cross-section of the basement lamina of a capillary is

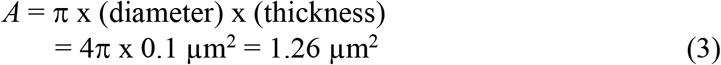

Speed of flow along basement lamina =

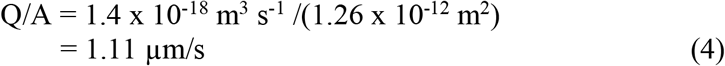

The data in Fig. 2C show that for frequencies in the range of the mouse heart beat (7 - 12 Hz; [53] and a pressure gradient of 40 cm H_2_O over 0.4 cm, dx/dt in the microburette ≈ 40 μm/s. Adjusting for the different diameter, the speed of flow in the agar column = 40 × (0.38/1)^2^ μm/s = 5.78 μm/s. Making the approximation that Darcy’s Law applies, so speed ∝ pressure gradient ([32], the pressure gradient needed for the speed through the agar gel to equal that along the basement lamina (Eq. (4)) is

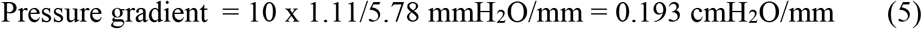

Little is known about pressure gradients in the brain. Holter et al. [21] considered that the upper possible limit was 1.36 cmH_2_O/mm, which is more than seven times the estimate in Equ. (5), and so could drive the hypothetical flow with a large margin.

More relevant is the pressure drop not in the neural tissue but between the CSF in the periarteriolar space at the upstream end of a capillary and the perivenular space at the downstream end: for a capillary with a typical length of 100 μm [2] the requirement would be 0.0193 cmH_2_O (Eq. 5). The pressure drop along the *inside* of a capillary can be as high as 34 cmH_2_O [69] which is 1761 times as big. Hence, sufficent energy to compress the basement lamina is available in the vicinity.

We conclude that although we do not know whether the effect of pulsation on the basement lamina causes greater or less enhancement of flow than the solenoid in our model, it would be rash to dismiss old [14] and recent [16–18] suggestions that CSF flows through a paracapillary pathway. In our model system, the transport increased with the frequency of pulsation over the range 0-5 Hz (Fig. 2C), and with the depth of compression, up to at least 1/6 of the external diameter of the gel-filled tube (Fig. 2B). An analogous progressive effect on the capillary basement lamina might account for the beneficial effect of physical exercise that involves increased cardiac output. This would complement other conditions that favour CSF distribution, such as slow-wave sleep [70].

## Supporting information

S1. Detailed methods

S2. Pulsation reduces transport of a solute through an open flexible tube

## Supporting Information

**S1. Detailed methods for the gel-filled tubes.**

**S2. Pulsation reduces transport of a solute through an open flexible tube.**

## Data accessibility

The code used to control the solenoid is available at https://github.com/GolfPapaEcho/CSF.

## Authors’ contributions

MH wrote the code for the microcomputers, analysed data, plotted graphs and purchased material. JAC did the experiments, analysed data, drafted figures and wrote the text. Both authors gave final approval for publication and agree to be held accountable for the work.

## Acknowledgements

The authors thank David Crofts for gifts of electronic components.

