## Supplementary material for "Intermittent compression increases flow through a gel: a mechanical model perhaps relevant to flow of cerebrospinal fluid": S1. Detailed methods

**S1. Detailed methods for the gel-filled tubes.**

Gel columns.

400 mg of agar agar (Top Op Foods Ltd, Stanmore, UK), was added to 40 ml of 0.15 M NaCl in a beaker which was warmed in a water bath at about 90 ºC. The mixture was stirred for 30 min. A10 ml syringe with a 19G needle, shortened and blunted with a diamond file, was used to inject columns of the warm agar into several 40 - 80 mm lengths of the silicone tubing i.d. 1.0 mm, o.d. 3.0 mm. The filled tubes were then stored in 0.15M NaCl at room temperature for up to a week.

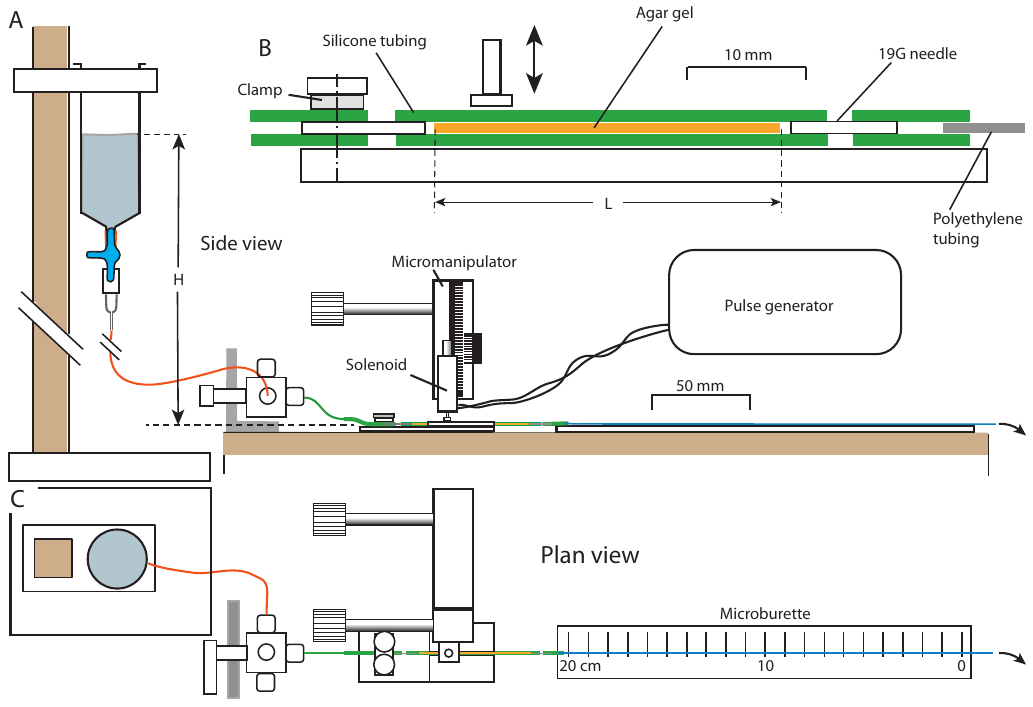

**Fig S1. The apparatus for the effect of pulsation**. A side view, C plan view. B Enlargement showing the gel column under the solenoid and the connections to the inflow and the microburette.

Apparatus for flow measurement

As shown in Fig S1A, to apply a longitudinal pressure gradient to the gel-filled tube, a filter stand was used to support a reservoir (a 50 ml syringe) containing 0.15 M NaCl with its surface at a height H above above the horizontal gel-filled tube. H was set at 200, 300, 400 or 500 mm. The fluid was led through a four-way distribution valve (HVP 86788, Hamilton, Bonaduz, CH) to 1 mm i.d. silicone tubing then the gel column, then a microburette consisting of low density polyethylene tubing (Portex, Smith Medical) with a nominal internal diameter of 0.38 mm, held with sellotape along 200 mm of a ruler scale.

Mounting the gel column

The silicone tube containing gel was cut to 10 mm longer than the desired length, *L*, of column. The 29G needle of an insulin syringe was cut to 5 mm with a diamond file and filled with 0.15M NaCl. It was inserted in each end of a length of tube and gel was flushed out and replaced by the saline solution. The tube from the reservoir ended in a length of 19G needle (Fig S1B); saline was allowed to flow until the needle was full; then the tip of the needle was inserted in one end of the gel tube. To avoid pressure on the gel, the distribution valve was turned to an open port and the needle further inserted to about 4 mm. Similarly, a length of 19G needle was inserted in the downstream end of the gel tube. This led to another length of 1 mm silicone tubing, which was filled with saline, and brought up to the microburette tubing, which was also full. When the microburette tube was pushed into the silicone tube, a bubble invariably entered it and was driven a short way down the scale. The advancing position of the downstream end of this bubble, best seen with torchlight and a loupe, was used to measure the flow. The timing between measurements was chosen so that the displacement was at least 1 mm: for overnight runs it could be more than 50 mm.

The solenoid

A 5 V solenoid (The Pi Hut, SKU:ADA2776) with a piston diameter of 3.5 mm was held vertically above the gel tube on a micromanipulator, about 10 mm from its upstream end. Its vertical position was determined to 0.1 mm with a Vernier scale. It's current was switched by a transistor (BFY51) controlled by a Raspberry Pi zero microcomputer. Although the duty cycle could be varied, it was kept at 50%. The Raspberry Pi was controlled by wifi from a MacBook Pro using the program 'Solenoid.py' on https://github.com/GolfPapaEcho/CSF.
