## Supplementary material for "Intermittent compression increases flow through a gel: a mechanical model perhaps relevant to flow of cerebrospinal fluid": S2. Pulsation reduces transport of a solute through an open flexible tube

S2. Pulsation reduces transport of a solute through an open flexible tube.
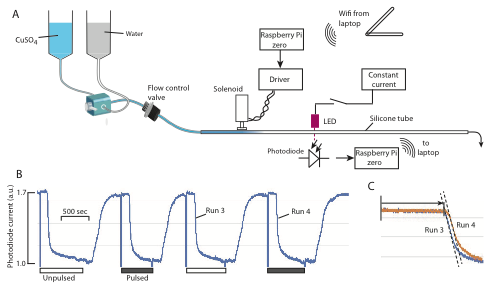

Fig S2. **A** Scheme for measuring the transport of a solute along an elastic tube. The photodiode was about 20 cm downstream of the two-way valve. Flow control valve: Biorad FCV. **B** When the solution was switched from water to CuSO_4_ solution (bars) the photodiode current fell after a delay. Filled bars indicate that the solenoid was activated at 5 Hz, 50% duty cycle. **C** Superimposed traces of two runs in **B** illustrating how, on average, the transport of CuSO_4_ was delayed when the tube was pulsed.
